# Cell size distribution of lineage data: analytic results and parameter inference

**DOI:** 10.1101/2020.12.24.424287

**Authors:** Chen Jia, Abhyudai Singh, Ramon Grima

## Abstract

Recent advances in single-cell technologies have enabled time-resolved measurements of the cell size over several cell cycles. This data encodes information on how cells correct size aberrations so that they do not grow abnormally large or small. Here we formulate a piecewise deterministic Markov model describing the evolution of the cell size over many generations, for all three cell size homeostasis strategies (timer, sizer, and adder). The model is solved to obtain an analytical expression for the non-Gaussian cell size distribution in a cell lineage; the theory is used to understand how the shape of the distribution is influenced by the parameters controlling the dynamics of the cell cycle and by the choice of cell tracking protocol. The theoretical cell size distribution is found to provide an excellent match to the experimental cell size distribution of *E. coli* lineage data collected under various growth conditions.

## Introduction

Cell size plays an important role in cellular processes; e.g. changes in cell volume or surface area have profound effects on metabolic flux and nutrient exchange [1], and therefore it stands to reason that cell size should be actively maintained. In order for cells to achieve and maintain some characteristic size (size homeostasis), the amount of growth produced during the cell cycle must be controlled such that, on average, large cells at birth grow less than small ones.

There are three popular phenomenological models of cell size control leading to size homeostasis [2]: (i) the timer strategy which implies a constant time between successive divisions; (ii) the sizer strategy which implies cell division upon attainment of a critical size, and (iii) the adder strategy which implies a constant size addition between consecutive generations. The timer strategy is not viable for exponentially growing cells; in this case, size fluctuations diverge as the square root of the number of consecutive cell divisions implying that the timer strategy cannot maintain stable size distributions [3]. In contrast, if cells grow linearly, a timer strategy is viable as a means to maintain size homeostasis [4]. Several studies have proposed that the sizer and adder strategies can explain experimental data in bacteria, yeast, and mammalian cells [5–10]. Cell-size control mechanisms likely vary depending on growth conditions, strains, and species; for instance in *Escherichia coli* (*E. coli*), evidence suggests a sizer mechanism in slow growth conditions and an adder in fast growth conditions [11].

Cell size statistics can be computed using data from cell lineages or population snapshots. To observe a single cell lineage, at each cell division event, one keeps track of only one of the newborn cells (daughter cells); thus at an arbitrary time point, only a single cell is observed. Whereas to observe population snapshots, one tracks both daughters of each mother cell in the population and thus the evolution of the whole population over time. Recently, mathematical models have shown that cell size statistics calculated using lineage data, e.g. collected using mother machines, can vary considerably from those obtained from population snapshot data, e.g. collected using flow cytometry [12–14]. In fact, differences between these two types of measurements are also observable in protein and mRNA count statistics [15–17].

Modelling has elucidated various other interesting insights into cell size statistics, however to our knowledge no study thus far has attempted to explain the complex shapes of cell size distributions computed from many generations of cell lineage measurements. This is because such high throughput data has become available only recently [18] and also since the majority of modelling approaches have analytically derived expressions for the first few moments of cell size statistics — these are not enough to characterize the highly non-Gaussian distributions of cell size computed from a cell lineage (see Fig. 1(a) for a typical distribution for an *E. coli* lineage). These histograms are characterized by three features: a fast increase in the size count for small cells, a slow decay in the size count for moderately large cells, and a fast decay in the size count for large cells. We note that these distributions contain much more information than birth size distributions previously derived [19], since they reflect the full cell cycle dynamics.

**Fig. 1.**
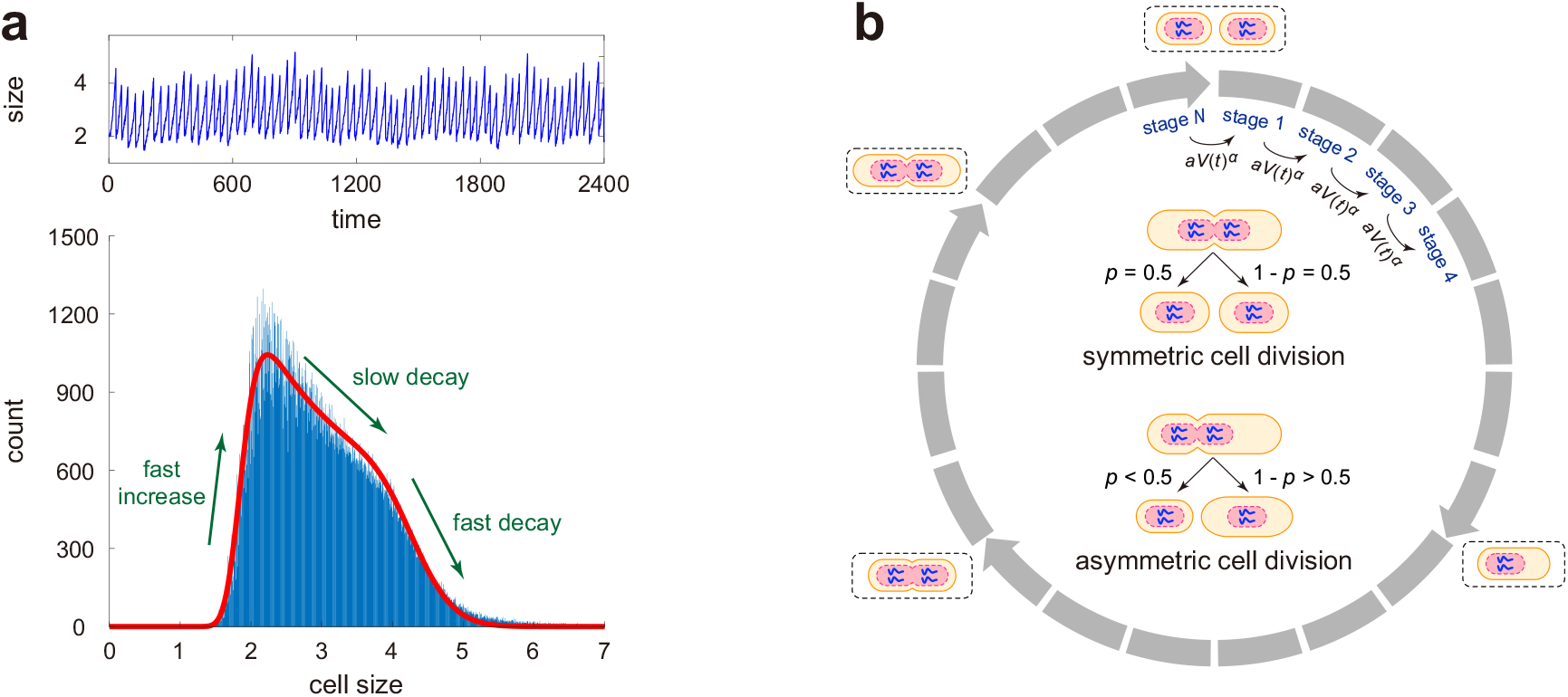
Cell size dynamics and a stochastic model describing it. (**a**) Single-cell time course data of cell length along a typical cell lineage measured in *E. coli* at 37°C (upper) and the histogram of cell sizes along all cell lineages (lower). The data shown are published in [7]. The cell size distribution computed from cell lineage measurements has an uncommon shape that is characterized by three features: a fast increase in the size count for small cells, followed by a slow decay for moderately large cells and a fast decay for large cells. (**b**) Schematic illustrating a detailed model of cell size dynamics describing cell growth, multiple effective cell cycle stages, cell-size control, and symmetric or asymmetric partitioning at cell division (see inset graph). Each cell can exist in *N* effective cell cycle stages. The transition rate from one stage to the next at a particular time *t* is proportional to the *α*th power of the cell size *V*(*t*) with *α* > 0 being the strength of cell-size control and *a* > 0 being the proportionality constant. This guarantees that larger cells at birth divide faster than smaller ones to achieve size homeostasis. At stage *N*, a mother cell divides into two daughters that are typically different in size via asymmetric cell division. Symmetric division is the special case where daughters are equisized.

Here we develop a complete analytical theory of the cell size distribution in cell lineages. We formulate and solve a piecewise deterministic Markov model describing the evolution of the cell size over many generations, for all three size homeostasis strategies (timer, sizer, and adder). The model takes into account the major features responsible for the underlying dynamics: cell birth following division (including the asymmetric case and partitioning noise), exponential cell growth (including the case of noisy growth rates), variability in the duration of the cell cycle, and the user-defined choice of single-cell tracking protocols when division occurs, e.g. tracking always the smaller daughters, tracking always the larger daughters, or randomly picking one of the two daughters. The analytical solutions for the cell size distribution enable us to understand how the highly non-Gaussian shape of the distribution emerges from the underlying biophysical processes. Finally by matching the analytical to the experimental cell size and doubling time distributions, we infer the values of various model parameters in *E. coli* for three different growth conditions.

## Results

### Model specification

Here we consider a detailed model of cell size dynamics across the cell cycle which is similar to the model proposed in [20] but has more complicated cell division mechanisms such as asymmetric and stochastic partitioning (see Fig. 1(b) for an illustration). The model is based on a number of assumptions that are closely tied to experimental data. The assumptions are as follows.

1. The size of each cell grows exponentially in each generation with growth rate *g*. This assumption is supported by experiments in many cell types [21].
2. Each cell can exist in *N* effective cell cycle stages, denoted by 1, 2,…, *N*. The transition rate from one stage to the next at a particular time is proportional to the *α*th power of the cell size at that time, with *a* > 0 being the proportionality constant [20]. In other words, the transition rate between stages at time *t* is equal to *aV*(*t*)^*α*^, where *α* > 0 is the strength of cell-size control and *V*(*t*) is the cell size at that time. Under this assumption, larger cells at birth have larger transition rates between stages and thus, on average, have lesser cell cycle duration and lesser volume change than smaller ones; in the way size homeostasis is achieved. Examples of possible biophysical mechanisms that can explain the power law form of the transition rate have been discussed in [20]. Let *V_b_* and *V_d_* denote the cell sizes at birth and at division in a particular generation, respectively. Then the increment in the *α*th power of the cell size across the cell cycle, 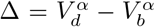, has an Erlang distribution with shape parameter *N* and mean *A* = *Nαg/a* (see Supplementary Section 1 for the proof). The quantity Δ will be referred to as the generalized added size in what follows. In our model, the noise in the generalized added size, characterized by the coefficient of variation squared, is equal to 1/*N*. As *N* increases, the generalized added size, as well as *V_b_* and *V_d_* themselves, have smaller fluctuations. Since the cell cycle duration is given by *T* = (1/*g*) log(*V_d_/V_b_*), an increasing *N* also results in lesser fluctuations in the doubling time. Hence, our model allows the investigation of the influence of cell cycle duration variability on cell size dynamics. We next focus on three crucial special cases. When *α* → 0, the transition rate between stages is a constant and thus the doubling time has an Erlang distribution that is independent of the birth size; this corresponds to the timer strategy. When *α* = 1, the added size *V_d_* – *V_b_* has an Erlang distribution that is independent of the birth size; this corresponds to the adder strategy. When *α* → ∞, the *α*th power of the cell size at division, 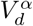, has an Erlang distribution that is independent of the birth size; this corresponds to the sizer strategy. Intermediate strategies are naturally obtained for intermediate values of *α*; timer-like control is obtained when 0 < *α* < 1 and sizer-like control is obtained when 1 < *α* < ∞ [20].
3. Cell division occurs when the cell transitions from effective stage *N* to the next stage 1. At division, most previous papers assume that the mother cell divides into two daughters that are exactly the same in size via symmetric partitioning [19, 22–25]; however, asymmetric cell division is common in biology. For instance, *Saccharomyces cerevisiae* divides asymmetrically into two daughters with different sizes. *Escherichia coli* may also undergo asymmetric division with old daughters receiving fewer gene products than new daughters [26]. Here we follow the methodology that we devised in [27] and extend previous models by considering asymmetric partitioning at cell division: the mother cell divides into two daughters with different sizes.

If the partitioning of the cell size is symmetric, we track one of the two daughters randomly after division [28, 29]; if the partitioning is asymmetric, we either track the smaller daughter or track the larger daughter after division [30, 31]. Hence our model corresponds to cell lineage measurements performed using a mother machine. Let *V_d_* and 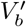 denote the cell sizes at division and just after division, respectively. If the partitioning is deterministic, then we have 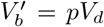, where 0 < *p* < 1 is a constant with *p* = 1/2 corresponding to the case of symmetric division, *p* < 1/2 corresponding to smaller daughter tracking, and *p* > 1/2 corresponding to larger daughter tracking. However, in naturally occurring systems, the partitioning is appreciably stochastic. In this case, we assume that the partition ratio 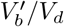 has a beta distribution with mean *p* [32], whose probability density function is given by

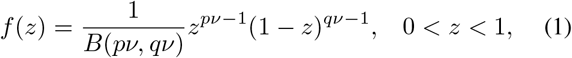

where *B* is the beta function, *q* = 1 – *p*, and *ν* > 0 is referred to as the sample size parameter. Then the change in the logarithm of the cell size at division, log 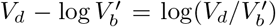, has the probability density function *μ*(*w*) = *e^−w^f*(*e^−w^*), which can be written more explicitly as

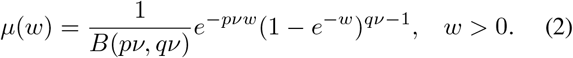

When *ν* → ∞, the variance of the beta distribution tends to zero and thus stochastic partitioning reduces to deterministic partitioning, i.e. *f*(*z*) = *δ*(*z* – *p*) and *μ*(*w*) = *δ*(*w* + log*p*).

We next describe our stochastic model of cell size dynamics across the cell cycle. The microstate of the cell can be represented by an ordered pair (*k, y*), where *k* is the cell cycle stage which is a discrete variable and *y* is the cell size which is a continuous variable. Let 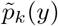 denote the probability density function of the cell size when the cell is in stage *k*. Note that the cell undergoes deterministic exponential growth in each stage and the system can hop between successive stages stochastically. Hence the evolution of the cell size dynamics can be described by a *piecewise deterministic Markov process* whose Kolmogorov backward equation is given by

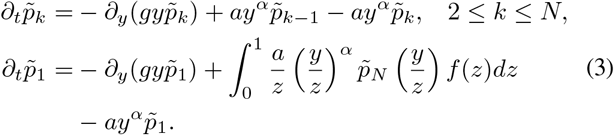

where *f*(*z*) is the function defined in Eq. (1). Similar hybrid models have, for example, been used to describe demographic noise in ecosystems [33] and single-cell stochastic gene expression [34, 35]. In the first equation above, the first term on the right-hand side represents the exponential growth of the cell size with growth rate *g*, the second and third terms represent the transition between stages whose transition rate is proportional to the *α*th power of the cell size *y*. In the second equation, the second term corresponds to the partitioning of the cell size at division.

To solve Eq. (3), the key step is to consider the dynamics of *the logarithmic cell size, x* = log *y*, rather than the original cell size *y*. This is because the dynamic equation for the former is easier to solve. Let *p_k_*(*x*) denote the probability density function of the logarithmic cell size when the cell is in stage *k*. Since the probability density functions of the original and logarithmic cell sizes are related by 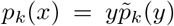, it follows from Eq. (3) that the evolution of the logarithmic cell size is governed by the following master equation:

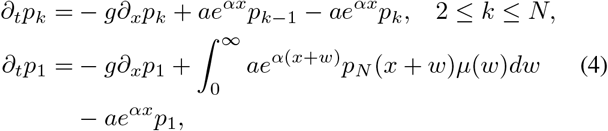

where *μ*(*w*) is the function defined in Eq. (2).

### Analytical distribution of the cell size along a cell lineage under deterministic partitioning

Recall that any probability distribution is fully determined by its characteristic function. Let 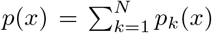 denote the probability density function of the logarithmic cell size. To obtain the analytical distribution of the cell size along a cell lineage, we introduce the characteristic function 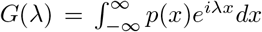, which is nothing but the inverse Fourier transform of *p*(*x*). For simplicity, we first focus on deterministic partitioning at cell division, i.e. *ν* → ∞. Despite the biological complexity described by our model, the characteristic function can still be solved exactly in steady-state conditions (see Supplementary Section 2 for the proof):

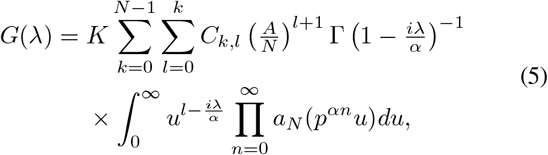

where *a_N_*(*u*) = (1 + *Au/N*)^−*N*^ is a function of *u* and

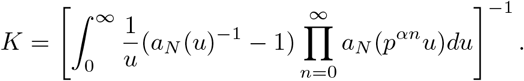

is a normalization constant. Since the Fourier transform and the inverse Fourier transform are inverses of each other, taking the Fourier transform of the characteristic function gives the steady-state probability density function *p*(*x*) of the logarithmic cell size. Finally, the probability density function of the original cell size *y* = *e^x^* along a cell lineage is given by

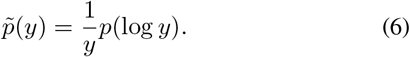

The analytical solution is ideal since it allows a fast exploration of large swathes of parameter space without performing stochastic simulations.

To gain deeper insights into the cell size distribution, we next consider the limiting case of *N* → ∞. In this case, the generalized added size △, as well as the cell cycle duration *T*, becomes deterministic and thus the system does not involve any stochasticity. As *N* → ∞, we have *a_n_*(*u*) = *e^−Au^* and thus the characteristic function can be simplified to a large extent as (see Supplementary Section 2 for the proof)

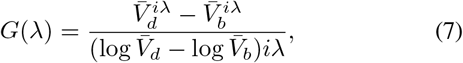

where

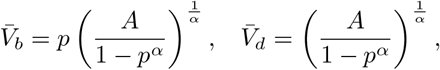

are two constants. Taking the Fourier transform of *G*(*λ*) shows that the logarithmic cell size has the uniform distribution

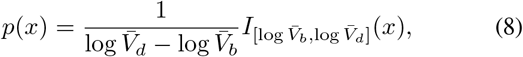

and thus the original cell size *y* = *e^x^* has the following distribution:

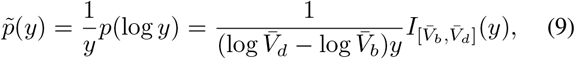

where *I_B_*(*x*) is the indicator function which takes the value of 1 when *x* ∈ *B* and the value of 0 otherwise. This indicates that when cell cycle duration variability is small, the cell size has a distribution that is concentrated on the finite interval 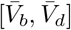, where 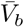 and 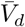 are the typical cell sizes at birth and at division, respectively.

Fig. 2(a)-(c) illustrate the distribution of the original cell size as a function of the parameters *N, α*, and *p*. It can be seen that as cell cycle duration variability become smaller (*N* increases), the analytical distribution given in Eq. (6) converges to the limit distribution given in Eq. (9). The cell size distribution has a regular shape for small *N*. As *N* increases, the shape of the distribution becomes more complicated. In particular, the distribution has three apparent sections: an exponential increase for small sizes, a power law decay for moderate sizes, and an exponential decay for large sizes. As *N* → ∞, the dynamics becomes deterministic and the distribution has a compact support, characterized by infinite slopes of the two shoulders. In addition, we find that the influence of *α* on the cell size distribution is similar to the influence of *N*. Finally, increasing *p* gives rise to a distribution that is more symmetric and more concentrated.

**Fig. 2.**
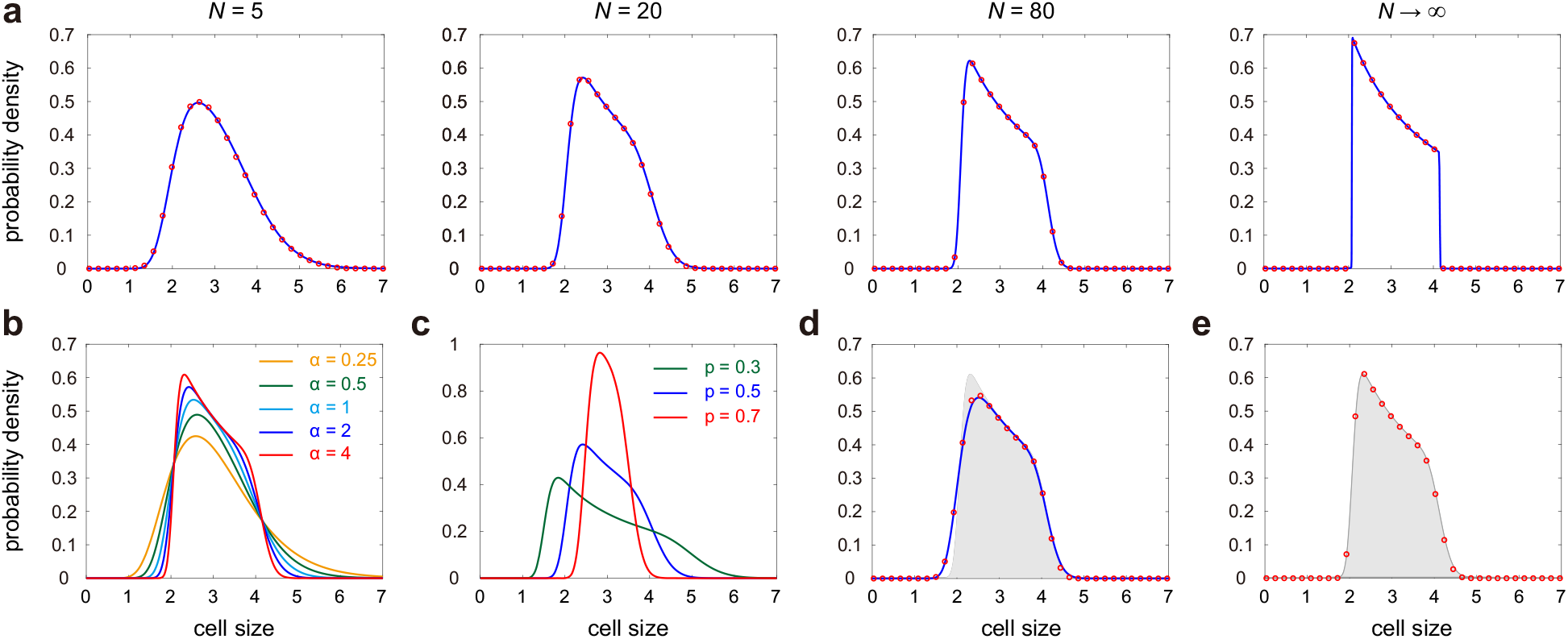
Influence of model parameters on the cell size distribution. (**a**) Cell size distribution as *N* increases. The red curve shows the analytical distribution given in Eq. (6) and the red circles show the distribution obtained using the stochastic simulation algorithm proposed in [25]. The parameters are chosen as *α* = 2, *p* = 0.5. (**b**) Cell size distribution as *α* varies. The parameters are chosen as *N* = 20, *p* = 0.5. (**c**) Cell size distribution as *p* varies. The parameters are chosen as *N* = 20, *α* = 2. (**d**) Comparison of the cell size distributions for the model with stochastic partitioning (blue curve and red circles) and the model with deterministic partitioning (solid grey region). The blue curve shows the approximate distribution given in Eq. (18) and the red circles show the distribution obtained from simulations. (**e**) Comparison of the cell size distributions for the model with stochastic growth rate (red circles) and the model with deterministic growth rate (solid grey region). In (d),(e), the parameters are chosen as *N* = 30, *α* = 3, *p* = 0.5. In (a)-(e), the growth rate is chosen as *g* = 0.02 and the parameters *A* and *a* are chosen so that 〈*V*〉 = 3 for the model with deterministic growth rate and deterministic partitioning. In (e) the standard deviation of the growth rate is 10% of the mean; here we assume that the growth rates for different generations are i.i.d. normally distributed random variables.

### Moments, noise, and skewness of the cell size distribution

Our analytic results can also be used to derive explicit expressions for several other quantities of interest. Recall that the probability density function *p*(*x*) for the logarithmic cell size and the probability density function 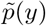 for the original cell size are related by Eq. (6). For any real number λ, the λth moment of the original cell size is given by

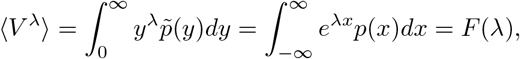

where *F*(λ) is the moment generating function of *p*(*x*). This shows that the λth moment of the original cell size is exactly the moment generating function of the logarithmic cell size taken value at λ. Since the moment generating function *F*(λ) and the characteristic function *G*(λ) are related by *G*(λ) = *F*(*i*λ), replacing the variable *i*λ in Eq. (5) by λ yields the moment generating function. Hence the λth moment of the original cell size is given by

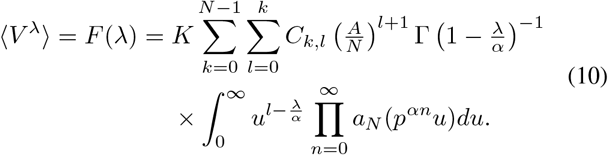

An important property of the cell size distribution is that it is a function of *A* = *Nαg/a*, which depends on the ratio of *g* and *a*. Therefore, different growth rates g may lead to the same size distribution whenever *g/a* is kept constant. In single-cell experiments, the noise in the cell size, characterized by the coefficient of variation squared, is given by

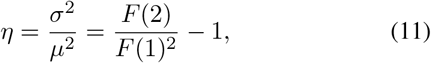

where *μ* is the mean and *σ*^2^ is the variance. Fig. 3(a),(b) illustrate the noise *η* as a function of *N*, α, and *p*. Clearly, the fluctuations in the cell size become smaller with the increase of all the three parameters (see also Fig. 2). This implies that small cell cycle duration variability and sizer-like strategy can lead to a more accurate control of the cell size.

**Fig. 3.**
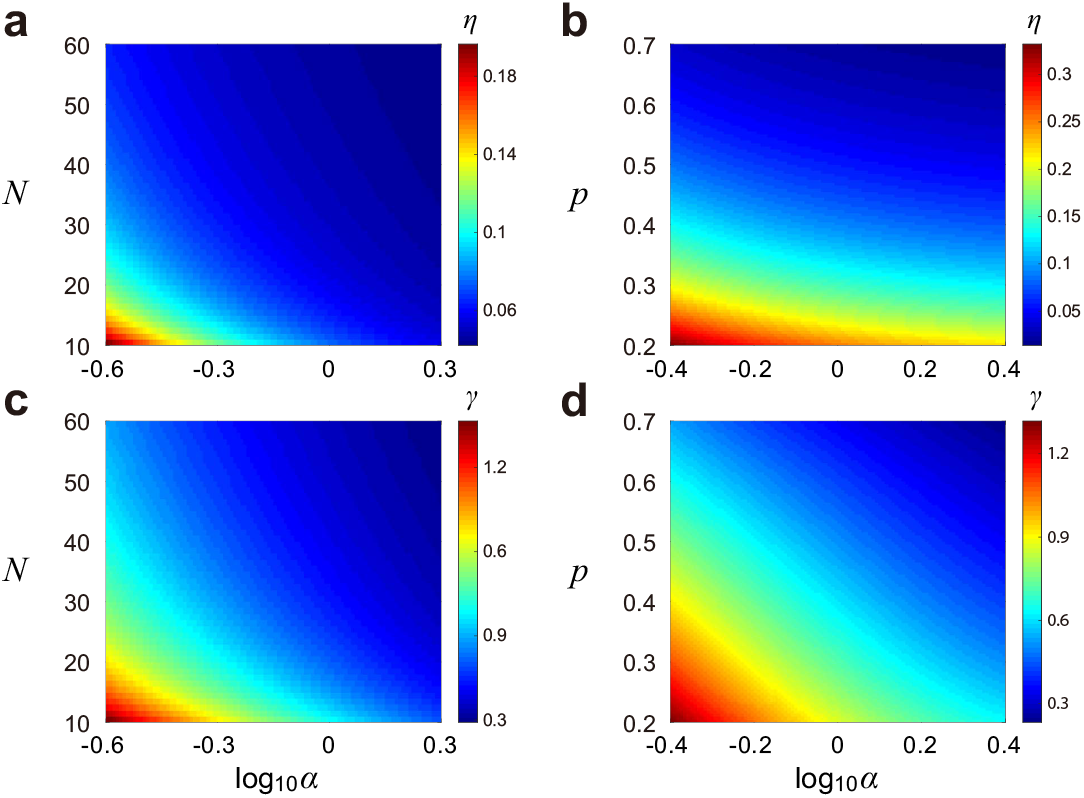
Noise and skewness of the cell size distribution. (**a**) Heat map of the noise *η* versus *α* and *N*. (**b**) Heat map of the noise *η* versus *α* and *p*. (**c**) Heat map of the skewness *γ* versus *α* and *N*. (**d**) Heat map of the skewness *γ* versus *α* and *p*. The parameters are chosen as *p* = 0.5 in (a),(c) and *N* = 20 in (b),(d). In (a)-(d), the parameter *A* is chosen so that the mean cell size 〈*V*〉 = 3.

A special case occurs when the cell cycle duration variability is very small, i.e. *N* ≫ 1. In this case, replacing the variable iλ in the characteristic function Eq. (7) by λ yields

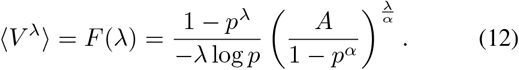

Thus the noise in the cell size is given by

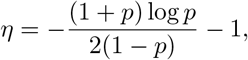

which is a decreasing function of *p*. Note that when *N* is small, the noise *η* is a function of both *α* and *p* (Fig. 3(b)). However, when *N* is large, the noise only depends on *p*. It is easy to see that the noise in the cell size tends to infinity as *p* → 0 and tends to zero as *p* → 1. For the case of symmetric division (*p* = 0.5), the noise in the cell size is given by *η* ≈ 0.04, which shows that the standard deviation of the cell size is roughly 20% of the mean.

Recall that the skewness of the cell size distribution is defined as

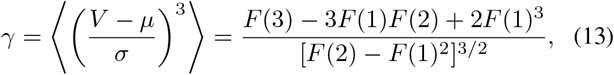

Fig. 3(c),(d) illustrate the skewness *γ* as a function of *N, α*, and *p*, from which we can see that the skewness increases with the decrease of all the three parameters. This implies that large cycle cycle duration variability, timer-like division strategy, and tracking the smaller daughter at division lead to larger skewness of the cell size distribution. Moreover, we find that the skewness is always positive, which means that the cell size distribution is always right-skewed. When *N* ≫ 1, it follows from Eq. (12) that the skewness only depends on *p* and is given by

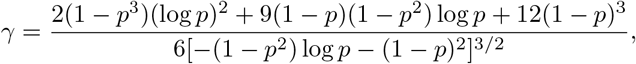

which is also a decreasing function of *p*.

### Analytical distribution of the cell cycle duration

In our model, the distribution of the doubling time can also be derived analytically in steady-state conditions. Actually, given that the birth size *V_b_* is known, the conditional probability density of the cell cycle duration *T* has been obtained in [20] as

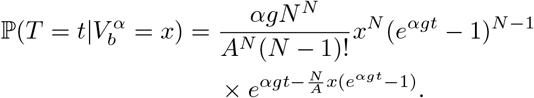

Here we compute the unconditional distribution of the cell cycle duration. To this end, we find that the Laplace transform of 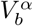 is given by (see Supplementary Section 6 for the proof)

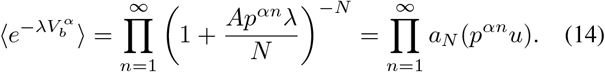

Taking the inverse Laplace transform gives the probability density function of 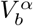. Finally, the distribution of the cell cycle duration *T* is given by

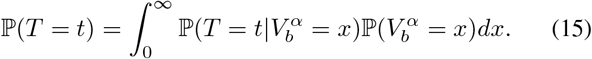

A special case occurs when *α* is large (strong cell-size control) or when *p* is small (smaller daughter tracking). Under the large *α* or small *p* approximation, the term *p^αn^* is negligible for *n* ≥ 2 and it suffices to keep only the first term in the infinite product given in Eq. (14). In this case, the inverse Laplace transform has an explicit expression and the birth size distribution is given by

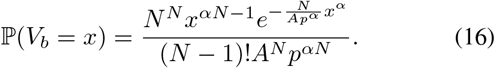

Inserting this equation into Eq. (15) yields the doubling time distribution

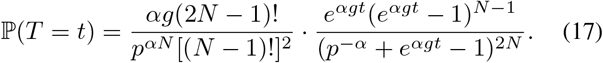

We emphasize that in the special case of *N* = 1, our model reduces to the model in [36] and the above two equations coincide with the results in that paper.

Recent experiments [17, 37–40] have shown that the cell cycle durations in various cell types are all well fitted by a gamma distribution. Therefore it is natural to ask whether the doubling time in our model shares the same property. To see this, we illustrate the doubling time distribution and its approximation by the gamma distribution as *N* and *α* vary (Fig. 4). It can be seen that the true distribution is in good agreement with its gamma approximation when *α* is small (Fig. 4(a),(b)). This is because a small *α* implies a timer-like size control, which leads to an approximately Erlang distributed doubling time due to the effect of multiple cell cycle stages and constant transition rates between them. When *α* is large, there are some slight differences between them for small *N* (Fig. 4(c)); compared with the gamma approximation, the true distribution is more symmetric around its mean. However, when *N* is large, they are very close to each other and both well fitted by a normal distribution (any gamma distribution converges to the normal distribution as the shape parameter tends to infinity, see Fig. 4(d)).

**Fig. 4.**
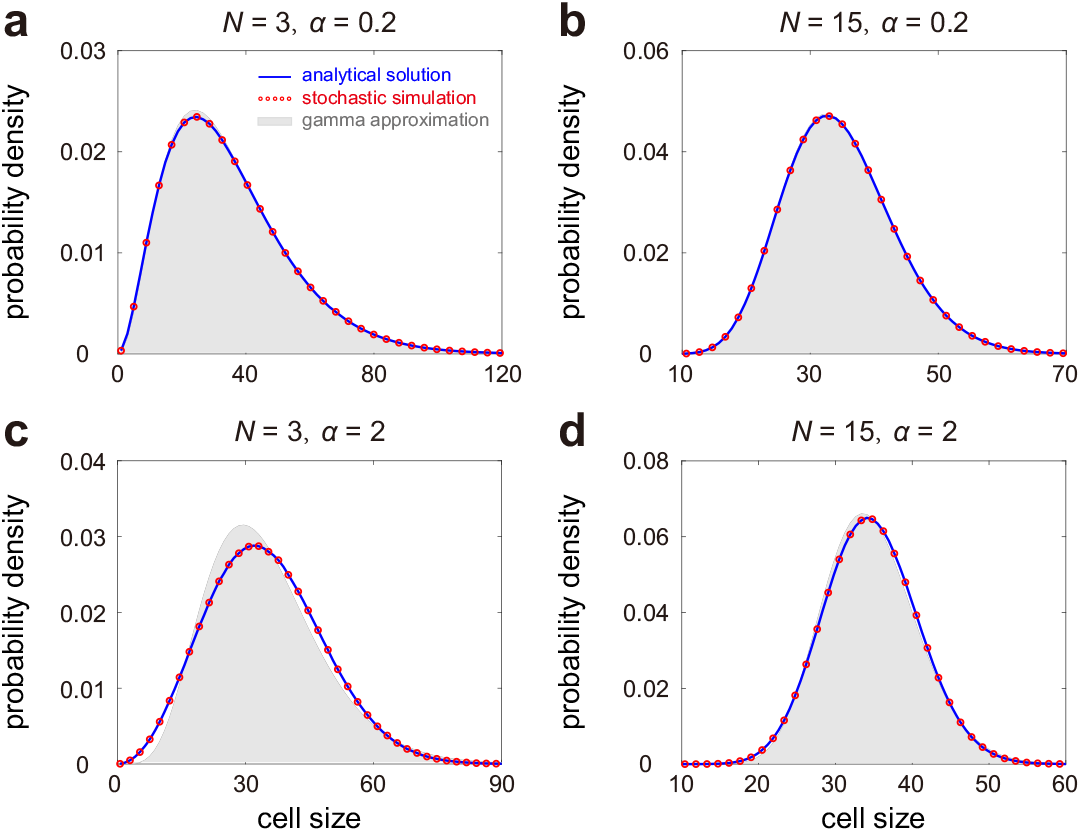
Distribution of the cell cycle duration and its approximation by the gamma distribution. We use the information of the sample mean and sample variance of the true distribution to determine the two parameters involved in the gamma approximation. (**a**) Large cell cycle duration variability and small size control strength. (**b**) Small cell cycle duration variability and small size control strength. (**c**) Large cell cycle duration variability and large size control strength. (**d**) Small cell cycle duration variability and large size control strength. In (a)-(d), the blue curve represents the analytical distribution given in Eq. (15), the red circles represent the distribution obtained from simulations, and the grey region represents the gamma approximation. The parameters are chosen as *p* = 0.5, *g* = 0.02 and *A* and *a* are determined so that 〈*V*〉 = 3.

### Distribution of the cell size along a cell lineage under stochastic partitioning and stochastic growth rate

Thus far, the analytical distribution of the cell size is obtained when the partitioning at division is deterministic. In the presence of noise in partitioning, it is very difficult to obtain the explicit expression of the cell size distribution. Fortunately, in naturally occurring systems, the stochasticity in partitioning is often very small. For example, recent cell lineage data [7] suggested that the coefficient of variation of the partition ratio 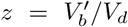 in *E. coli* is about 7% - 9%. When noise in partitioning is small, we obtain an approximate expression for the cell size distribution, whose moment generating function is given by (see Supplementary Section 3 for the proof)

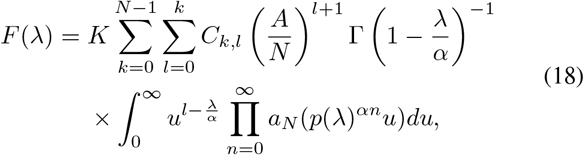

where *K* is a normalization constant and

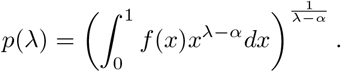

To see the effect of stochastic partitioning, we illustrate the cell size distributions under deterministic and stochastic partitioning in Fig. 2(d) with the standard deviation of the partition ratio *z* being 10% of the mean for the latter. Clearly, the approximate solution given in Eq. (18) matches the simulation results very well. In addition, it can be seen that noise in partitioning gives rise to larger fluctuations in the cell size, characterized by the smaller slope of the left shoulder of the cell size distribution.

In addition to noise in partitioning, there is another important source of stochasticity, i.e. noise in the growth rate *g*. In many biological systems, such noise is also very small. For example, recent cell lineage data [7] suggested that the coefficient of variation of the growth rate *g* in *E. coli* is about 7% - 8%. To see the influence of noise in the growth rate, we illustrate the cell size distributions under deterministic and stochastic growth rates in Fig. 2(e) with the standard deviation of *g* being 10% of the mean for the latter (here we assume that the growth rates for different generations are i.i.d. normally distributed random variables). Interestingly, we find that noise in the growth rate has very little effect on the cell size distribution; this is in sharp contrast to noise in partitioning which has an apparent effect on the cell size distribution.

### Random tracking protocol can lead to complex multimodal cell size distributions

If cell division is asymmetric, the two daughters are different in size and thus far we have assumed that the smaller/larger daughter (such as the bud/mother cell in budding yeast) is tracked after division [30, 31]. We have seen that whether the smaller or the larger daughter is tracked, the cell size distribution along a cell lineage is always unimodal and right-skewed, and larger daughter tracking yields lesser fluctuations in size than smaller daughter tracking. Next we consider another tracking protocol, namely where we track one of the two daughters randomly with probability 1/2 after division [7, 28, 29]. Clearly, the three types of tracking protocols (tracking a random daughter, the smaller daughter, or the larger daughter) are exactly the same for symmetric cell division; however, they are remarkably different for asymmetric cell division.

For the random tracking protocol, the probability density function of the partition ratio 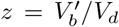 is given by (here the noise in partitioning is ignored)

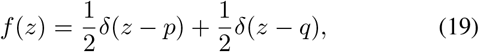

where 0 < *p* ≤ 1/2 is the ratio of the size of the smaller daughter to the size of the mother cell and *q* = 1 – *p*. Fig. 5 illustrates the simulated cell size distribution under the random tracking protocol. Interestingly, we find that the shape of the distribution undergoes two stochastic bifurcations as cell cycle duration variability becomes smaller (*N* increases). When *N* is small, the cell size distribution is in general unimodal (Fig. 5(a)), as in the case of smaller/larger daughter tracking. When *N* is moderate, random tracking is capable of producing a bimodal cell size distribution (Fig. 5(b)), where the two peaks can be attributed intuitively to the subpopulations of smaller and larger daughters, respectively. Surprisingly, when *N* is large, we find that random tracking can give rise to a complex cell size distribution that displays multiple peaks (Fig. 5(b)), two major peaks and some minor peaks. Increasing the cell cycle duration variability (decreasing *N*) smoothens the cell size distribution, by first removing the smaller peaks and then merging the two major peaks into one.

**Fig. 5.**
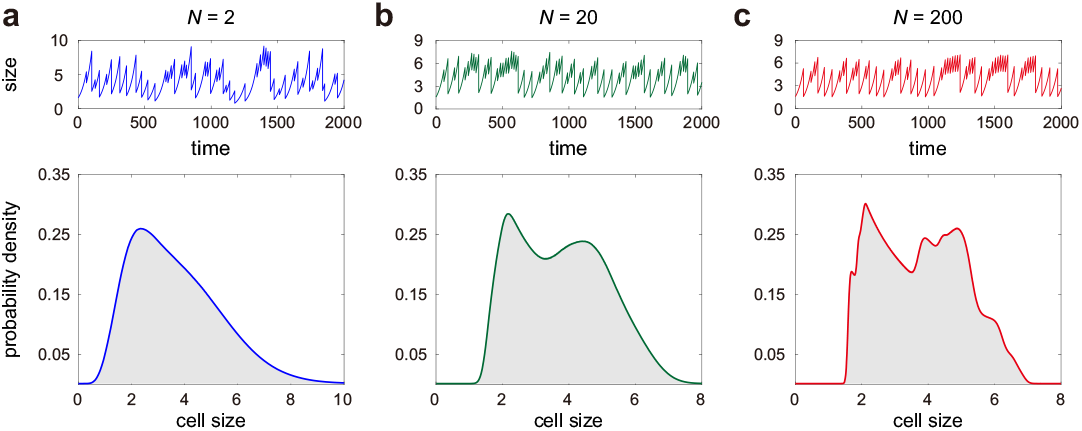
Cell size distribution for asymmetric cell division under the random tracking protocol. After division, one the two daughters is randomly tracked with probability 1/2. (**a**) Typical stochastic trajectory of the cell size (upper) and the cell size distribution (lower) in the case of large cell cycle duration variability (*N* = 2). (**b**) Same as (a) but for moderate cell cycle duration variability (*N* = 20). (**c**) Same as (a) but for small cell cycle duration variability (*N* = 200). In (a)-(c), the colored curve and the grey region show the cell size distributions obtained from two independently repeated stochastic simulations. The parameters are chosen as *p* = 0.3, *α* = 2, *A* = 25.

### Parameter inference using synthetic data

Recent breakthroughs in microfluidic devices have made it possible to monitor the single-cell volume dynamics along a cell lineage over many generations. Given such cell lineage data, an important question is whether all the parameters involved in our model can be inferred accurately. Parameter inference is crucial since it provides insights on the strength of cell-size control as well as cell cycle duration variability in various cell types.

The steps of our parameter estimation method are described as follows. First, the data of cell sizes at birth and at division in each generation, *V_b_* and *V_d_*, can be easily extracted from the cell lineage data. Since 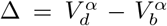 is Erlang distributed with shape parameter *N* and mean *A*, once the parameter *α* is determined, both the parameters *N* and *A* can be determined by fitting the data of 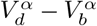 to an Erlang distribution. For clarity, let *N*(*α*) and *A*(*α*) denote the optimal estimates of *N* and *A* given the value of *α*. They can be inferred from the generalized added size △ as

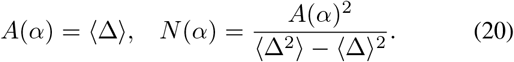

Next, the parameters *α* and *p* are determined by an optimal fit of the experimental to the theoretical cell size distribution using the least square criterion. Specifically, we determine *α* and *p* by solving the following optimization problem:

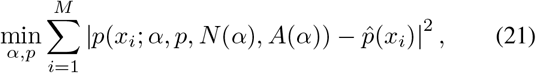

where *p*(*x*; *α,p,N,A*) is the theoretical cell size distribution given the parameters *α,p,N,A*, 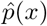 is the sample cell size distribution obtained from experiments, *x_i_* are some reference points, and *M* is the number of bins chosen. Once *α* and *p* are estimated, both *N* and *A* are automatically determined. The reason why we do not estimate *p* directly as the mean of the partition ratio 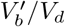 is that the cell size distribution is sensitive to the value of *p*. A comparatively small error in *p* will result in a comparatively large change in the cell size distribution.

Since the cell size distribution is a function of *A* = *Nαg/a*, which depends on the ratio of *g* and *a*, it is impossible to infer the growth rate *g* from the cell size distribution. Finally, the growth rate *g* is determined by an optimal fit of the experimental to the theoretical/simulated doubling time distribution using the least square criterion. Once *g* is inferred, the last parameter *a* can be determined from the estimated *α, N*, and *A* as *a* = *Nαg/A*.

**Table 1.** Parameter inference using synthesis data from the stochastic model. The cell lineage data are generated from the model with stochastic partitioning, where some noise is added to the growth rate *g* and the partition ratio *z* with the coefficients of variation of both parameters being chosen as 7%. For each set of model parameters, we generate synthetic data simulating 50 cell lineages. For each cell lineage, the model parameters are estimated by fitting the synthetic data to both the model with deterministic partitioning at cell division (Model I) and the model with stochastic partitioning (Model II). The value in each cell shows the mean and standard deviation of the estimated parameter computed over 50 cell lineages.

| | $p$ | $\alpha$ | $N$ | $A$ | $g$ | $a$ |
| --- | --- | --- | --- | --- | --- | --- |
| input parameters | 0.4 | 0.5 | 30 | 0.79 | 0.01 | 0.191 |
| estimated parameters using Model I | $0.40 \pm 0.0002$ | $0.44 \pm 0.02$ | $28.70 \pm 0.82$ | $0.65 \pm 0.06$ | $0.0100 \pm 0.0001$ | $0.195 \pm 0.015$ |
| estimated parameters using Model II | $0.40 \pm 0.0003$ | $0.50 \pm 0.04$ | $28.10 \pm 1.52$ | $0.79 \pm 0.11$ | $0.0100 \pm 0.0002$ | $0.179 \pm 0.016$ |
| | $p$ | $\alpha$ | $N$ | $A$ | $g$ | $a$ |
| input parameters | 0.5 | 1 | 20 | 2.08 | 0.02 | 0.192 |
| estimated parameters using Model I | $0.50 \pm 0.0003$ | $0.77 \pm 0.06$ | $21.15 \pm 0.88$ | $1.26 \pm 0.18$ | $0.0200 \pm 0.0002$ | $0.263 \pm 0.026$ |
| estimated parameters using Model II | $0.50 \pm 0.0003$ | $0.99 \pm 0.08$ | $19.20 \pm 1.14$ | $2.04 \pm 0.35$ | $0.0200 \pm 0.0003$ | $0.187 \pm 0.027$ |
| | $p$ | $\alpha$ | $N$ | $A$ | $g$ | $a$ |
| input parameters | 0.6 | 2 | 10 | 9.39 | 0.03 | 0.064 |
| estimated parameters using Model I | $0.60 \pm 0.0005$ | $1.29 \pm 0.04$ | $12.90 \pm 0.32$ | $2.73 \pm 0.20$ | $0.0301 \pm 0.0003$ | $0.183 \pm 0.010$ |
| estimated parameters using Model II | $0.60 \pm 0.0004$ | $1.92 \pm 0.04$ | $10.40 \pm 0.52$ | $8.29 \pm 0.57$ | $0.0300 \pm 0.0004$ | $0.072 \pm 0.004$ |

To verify the effectiveness of our method, we use our model to generating synthetic data of cell size dynamics. To make the time course data better mimic real biological process, we add some noise to both the growth rate *g* and the partition ratio *z*. We then perform parameter inference by fitting the noisy data to two models: the model with deterministic partitioning (Model I) and the model with stochastic partitioning (Model II). The parameters input to the synthetic data and the parameters estimated using the above method are given in Table 2, where three sets of input parameters are chosen to cover large swathes of parameter space and to include three types of control strategies (timer-like, adder, and sizer-like). It can be seen that fitting the noisy data to both models leads to an accurate estimation of *p* and *g*, and a relatively accurate estimation of *N*. However, fitting the data to Model I gives rise to a systematic underestimation of *α* and *A*, and an overestimation of *α* due to stochasticity in partitioning. Fitting the data to Model II can remarkably improve the accuracy of estimation of these three parameters.

**Table 2.**
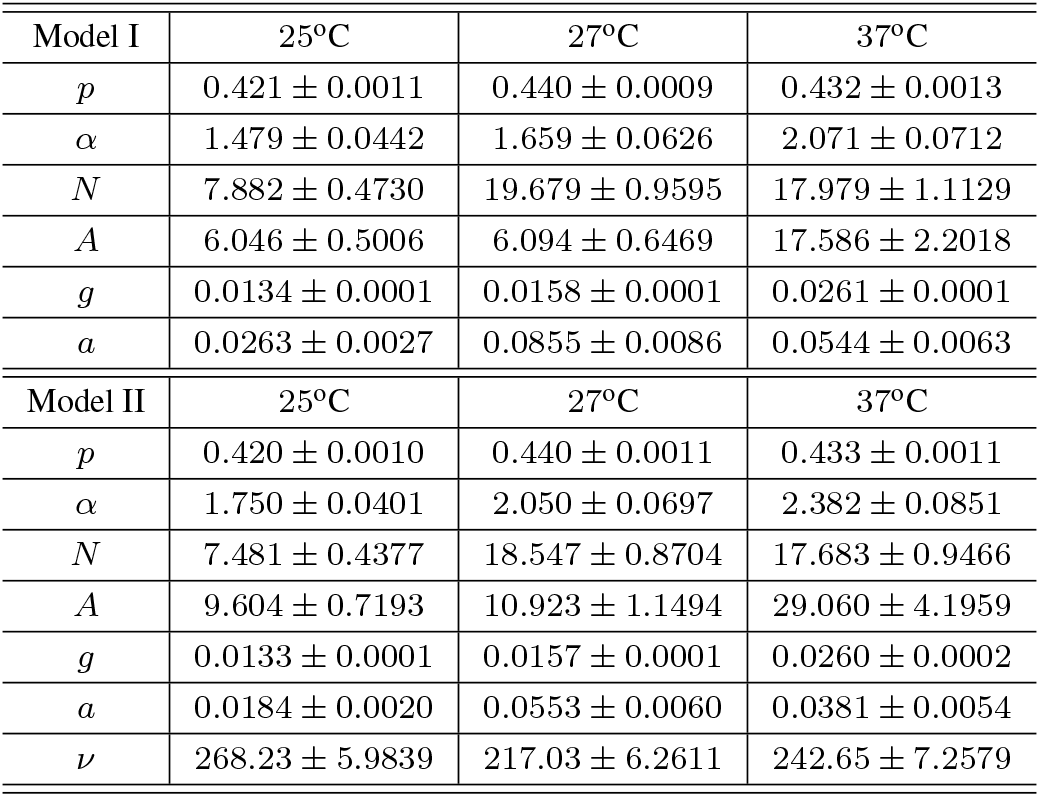
**The parameters estimated using cell lineage data** at three different temperatures. The parameters *p, α, N, A* are determined by fitting the experimental to the theoretical cell size distribution. The parameters *g* and *a* are determined by fitting the experimental to the theoretical doubling time distribution. Two theoretical models are used: the model with deterministic partitioning (Model I) and the model with stochastic partitioning (Model II). For model II, once the parameter *p* is estimated, the sample size parameter *v* in Eq. (1) can be inferred by fitting the partition ratio data to a beta distribution. The estimation error for each parameter was computed using bootstrap. Specifically, we performed parameter inference 50 times; for each estimation, the theoretical model was fitted to the data of 30 randomly selected cell lineages. The estimation error was then calculated as the standard deviation over 50 repeated samplings.

| Model I | 25°C | 27°C | 37°C |
| --- | --- | --- | --- |
| $p$ | $0.421 \pm 0.0011$ | $0.440 \pm 0.0009$ | $0.432 \pm 0.0013$ |
| $\alpha$ | $1.479 \pm 0.0442$ | $1.659 \pm 0.0626$ | $2.071 \pm 0.0712$ |
| $N$ | $7.882 \pm 0.4730$ | $19.679 \pm 0.9595$ | $17.979 \pm 1.1129$ |
| $A$ | $6.046 \pm 0.5006$ | $6.094 \pm 0.6469$ | $17.586 \pm 2.2018$ |
| $g$ | $0.0134 \pm 0.0001$ | $0.0158 \pm 0.0001$ | $0.0261 \pm 0.0001$ |
| $a$ | $0.0263 \pm 0.0027$ | $0.0855 \pm 0.0086$ | $0.0544 \pm 0.0063$ |
| Model II | 25°C | 27°C | 37°C |
| $p$ | $0.420 \pm 0.0010$ | $0.440 \pm 0.0011$ | $0.433 \pm 0.0011$ |
| $\alpha$ | $1.750 \pm 0.0401$ | $2.050 \pm 0.0697$ | $2.382 \pm 0.0851$ |
| $N$ | $7.481 \pm 0.4377$ | $18.547 \pm 0.8704$ | $17.683 \pm 0.9466$ |
| $A$ | $9.604 \pm 0.7193$ | $10.923 \pm 1.1494$ | $29.060 \pm 4.1959$ |
| $g$ | $0.0133 \pm 0.0001$ | $0.0157 \pm 0.0001$ | $0.0260 \pm 0.0002$ |
| $a$ | $0.0184 \pm 0.0020$ | $0.0553 \pm 0.0060$ | $0.0381 \pm 0.0054$ |
| $\nu$ | $268.23 \pm 5.9839$ | $217.03 \pm 6.2611$ | $242.65 \pm 7.2579$ |

### Experimental validation of the theory

To test our theory, we apply it to study the single-cell time course data of the cell size collected for *E. coli* in [18]. In this data set, the time course data of the cell length were recorded every minute for 279 cell lineages over 70 generations using a mother machine microfluidic device under three different growth conditions (25°C, 27°C, and 37°C). At the three temperatures, there are a total of 65, 54, and 160 cell lineages measured, respectively. Based on such data, it is possible to estimate all the parameters involved in our model at each temperature by fitting the data to both Model I and Model II. The estimated parameters and the estimation errors are listed in Table 2.

From the estimated parameters, it can be seen that both models lead to similar estimation of *p, N*, and *g*. However, the introduction of partitioning noise into the model leads to higher estimation of *α* and *A* and lower estimation of *a*; this is consistent with our observation for synthesis data. Regardless of the model used, the strength of cell-size control, *α*, is estimated to be 1.4 - 2.4 for all the three temperatures, implying that the size homeostasis strategy in *E. coli* is between the adder and the sizer. Moreover, higher temperature leads to a higher strength *α* than lower temperature.

Fig. 6(a),(b) illustrate the experimental cell size and cell cycle duration distributions using the data of all cell lineages versus the theoretical distributions using the estimated parameters. Here the theoretical distributions are plotted based on Model II but both models lead to similar distribution shapes. Interestingly, both experimental distributions at all the three temperatures coincide perfectly with our model, which implies that our model can indeed reproduce the cell size dynamics in *E. coli* very well. In addition, it verifies the main assumption of choosing the rate of moving from one effective cell cycle stage to the next to be a power law of the cell size.

**Fig. 6.**
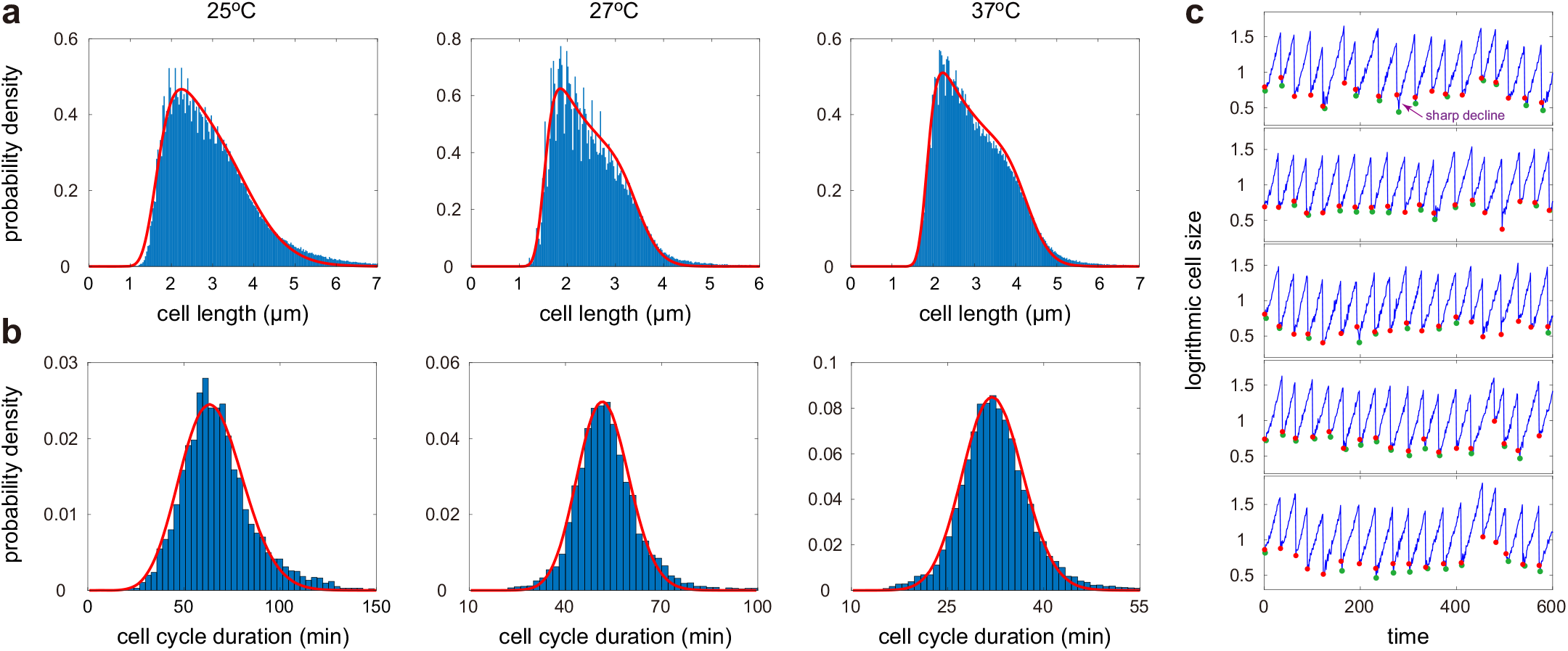
Fitting the experimental cell size and doubling time distributions to theory. (**a**) Experimental cell size distributions (blue bars) at the three temperatures and their optimal fitting to Model II (red curve) where partitioning is stochastic. Here the theoretical distribution is computed using Eq. (18). (**b**) Experimental doubling time distributions (blue bars) at the three temperatures and their optimal fitting to Model II (red curve), where the theoretical distribution is computed using stochastic simulations. (**c**) Five typical trajectories of cell size dynamics for cells at 37°C. The red dots show the cell sizes just after division and the green dots show the minimal cell sizes in each generation. For over 60% generations, there is a small abrupt decline in the cell size after division, shown as the sharp drop from a red dot to a green dot.

Typically, a mother cell divides into two daughters that are different in size due to stochasticity in partitioning and possible asymmetric cell division [32]. Note that the data of cell sizes just before division and just after division, *V_d_* and 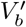, can be easily extracted from the cell lineage data and thus the parameter *p* can be estimated as the mean partition ratio 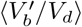. An interesting characteristic implied by the *E. coli* data is that at cell division, the smaller daughter is always tracked with the mean partition ratio *p* being estimated to be about 0.46 for all the three growth conditions (0.459 ± 0.040 for cells at 25°C, 0.461 ± 0.039 for cells at 27°C, and 0.464 ± 0.034 for cells at 37°C).

Recall that in our estimation procedure, we do not use the information of *V_d_* and 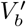 to determine the parameter *p*; rather, we infer *p* by an optimal fit of the experimental to the theoretical cell size distribution. The estimate of *p* in Table 2 is 0.42 - 0.44 for the three temperatures, which is slightly lower than the value of 0.46 estimated using *V_d_* and 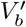. The reason of this discrepancy is that after cell division, over 60% generations have a small but sharp decline in the cell size (see Fig. 6(c) for the cell size dynamics of five typical cell lineages with the red dots being the cell sizes just after division and the green dots being the minimal cell sizes in each generation; the small sharp drop in the cell size after division is shown as the transition from a red dot to a green dot). Therefore, the realistic effective partition ratio should be computed using the green dots rather than the red dots. This explains why the parameter *p* estimated in Table 2 is lower than the mean partition ratio 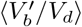.

## Summary and Discussion

In this work, we have analytically derived the cell size distribution of measurements obtained from a cell lineage. We have solved two models. The first model assumes that (i) the birth size is a fixed (generation independent) fraction of the division size in the last generation; (ii) the cell grows exponentially between birth and division events where the growth rate is a generation independent constant; (iii) the length of the cell cycle is stochastic; (iv) size homeostasis is enforced by timer-like, sizer-like, or adder strategies. A second model was also solved which relaxes the assumption (i) above, namely it allows for a stochastic ratio of the birth to division size.

The main features of the experimental cell size distribution in *E. coli*, namely a fast increase in the size count for small cells, a slow decay for moderately large cells, and a fast decay for large cells, are reproduced by the analytical solution of both models when the parameters *N* and *α* are large enough; this implies that these features emerge when the variability in the cell cycle duration is not too large and when adder or sizer-like mechanisms enforce size homeostasis. We also find that noise in partitioning at cell division (noise in the ratio of the birth to division size) has a considerable influence on the shape of the cell size distribution whereas noise in the growth rate hardly exerts any influence; this is in agreement with an earlier moment-based study [41].

Our theory predicts that large cell cycle duration variability, timer-like division strategy, and tracking the smaller daughter at division lead to larger skewness and coefficient of variation of the cell size distribution. We have furthermore shown that (i) the distribution of cell cycle duration that emerges from our model is well approximated by a gamma distribution that has been measured experimentally for many cell types [37]; (ii) if cells divide asymmetrically, they are tracked randomly after division, and cell cycle duration variability is intermediate or low, then the cell size distribution is multimodal.

Lastly, we have shown that the theoretical distributions provide an excellent fit to the experimental *E. coli* cell size and doubling time distributions reported in [18] for three different growth conditions. This match provides support for the implicit assumption of our model that the speed of the cell cycle (the transition rate between effective stages) monotonically increases with the cell volume and specifically has a power law dependence on the cell volume. We note that whilst this law may be compatible with certain biophysical mechanisms [25], more likely it is simply a phenomenological means to model cell size homeostasis; in fact more generally and beyond the context of our model, the usage of kinetic rates with power laws has found widespread applications in the effective modelling of complex biochemical kinetics in cells [42]. Finally based on the matching of the experimental to the theoretical cell size and doubling time distributions, we have estimated all the model parameters directly from *E. coli* cell lineage data and found that the strength *α* of cell-size control exhibits a weak increase with temperature. The estimated values of *α* (using Model II, the most accurate model in this paper) ranging between 1.8 and 2.4 confirm the results of a previous analysis [7] that neither an adder mechanism (*α* = 1) nor a sizer mechanism (*α* ≫ 1) can completely account for cell size homeostasis.

Concluding, the major advance in our work is the analytic derivation of the cell size distribution of lineage measurements, whereas previous studies focused more on population measurements or moments of lineage measurements. The advantages of the analytical distribution are (i) the ease and speed with which one can explore the dependence of cell size statistics on parameters across large swathes of parameter space (compared to stochastic simulations); (ii) the reliable estimation of parameters from data based on distribution matching which is generally much more robust than moment-based estimation [43]. The present model presents a framework onto which one can build further biological realism; current research work aims to extend the model to include gene expression and its correlation to cell size resulting in concentration homeostasis of mRNAs and proteins [2, 44–48].

## Acknowledgements

C. J. acknowledges support from the NSAF grant in National Natural Science Foundation of China with grant No. U1930402. A. S. is supported by the National Institute of Health Grant 1R01GM126557. R. G. acknowledges support from the Leverhulme Trust (RPG-2018-423).

## Competing interests

The authors declare that they have no competing interests.

## Data and materials availability

All data needed to evaluate the conclusions in the paper are present in the paper and in [18].

## Notes

### Competing Interest Statement

The authors have declared no competing interest.

