## Supplemental Information for "Cell size distribution of lineage data: analytic results and parameter inference"

### Supplemental Material for Cell size distribution of lineage data: analytic results and parameter inference

Chen Jia[1], Abhyudai Singh[2], Ramon Grima[3,*]
[1]Applied and Computational Mathematics Division, Beijing Computational Science Research Center, Beijing, China
[2]Department of Electrical and Computer Engineering, University of Delaware, Newark, Delaware, U.S.A.
[3] School of Biological Sciences, University of Edinburgh, U.K.

#### 1 Distribution of the generalized added size

Let $V_b$ and $V_d$ denote the cell sizes at birth and at division in a particular generation, respectively. In the main text, we have stated that the generalized added size $V_d^\alpha - V_b^\alpha$ is Erlang distributed with shape parameter $N$ and mean $A$, where $A = N\alpha g/a$. To see this, note that when $V_b$ is fixed, the cell size in this generation is given by $V(t) = V_be^{gt}$. Since the transition rate from one stage to the next at time $t$ is equal to $aV(t)^\alpha$, the distribution of the transition time $T$ is given by

$$\mathbb{P}(T > t) = e^{-\int_0^t aV(s)^\alpha ds} = e^{-\int_0^t aV_b^\alpha e^{\alpha gs}ds} = e^{-\frac{aV_b^\alpha}{\alpha g}(e^{\alpha gt}-1)}. \tag{1}$$

This shows that

$$\mathbb{P}(V_b^\alpha(e^{\alpha gT} - 1) > t) = e^{-\frac{at}{\alpha g}}.$$

Hence $V_b^\alpha(e^{\alpha gT} - 1)$ is exponentially distributed with mean $\alpha g/a$. Note that $V_b^\alpha(e^{\alpha gT} - 1)$ is the increment of the $\alpha$th power of the cell size in a particular cell cycle stage. Therefore, the total increment of the $\alpha$th power of the cell size in each generation is the independent sum of $N$ exponentially distributed random variables with mean $\alpha g/a$. This shows that $V_d^\alpha - V_b^\alpha$ has an Erlang distribution with shape parameter $N$ and mean $N\alpha g/a = A$.

#### 2 Cell size distribution under deterministic partitioning

##### 2.1 General case

Here we compute the analytical distribution of the cell size. For simplicity, we first focus on the case of deterministic partitioning at cell division, i.e. $\mu(y) = \delta(y + \log p)$. In this case, Eq. (4) in the main text reduces to

$$\begin{aligned}
\partial_t p_k &= - g\partial_x p_k + ae^{\alpha x}p_{k-1} - ae^{\alpha x}p_k, \quad 2 \le k \le N, \\
\partial_t p_1 &= - g\partial_x p_1 + ap^{-\alpha}e^{\alpha x}p_N(x - \log p) - ae^{\alpha x}p_1.
\end{aligned} \tag{2}$$

To proceed, for each cell cycle stage $k$, we introduce the moment generating function

$$F_k(\lambda) = \int_{-\infty}^{\infty} p_k(x)e^{\lambda x}dx, \quad F(\lambda) = \int_{-\infty}^{\infty} p(x)e^{\lambda x}dx,$$

where $p(x) = \sum_{k=1}^{N} p_k(x)$ is the probability density function of the logarithmic cell size. Then Eq. (2) can be converted to the following differential equations:

$$\begin{aligned} \partial_t F_k(\lambda) &= g\lambda F_k(\lambda) - aF_k(\lambda+\alpha) + aF_{k-1}(\lambda+\alpha), \quad 2 \le k \le N, \\ \partial_t F_1(\lambda) &= g\lambda F_1(\lambda) - aF_1(\lambda+\alpha) + ap^{\lambda} F_N(\lambda+\alpha). \end{aligned}$$

At the steady state, we have

$$\begin{aligned} &g\lambda F_k(\lambda) - aF_k(\lambda+\alpha) + aF_{k-1}(\lambda+\alpha) = 0, \quad 2 \le k \le N, \\ &g\lambda F_1(\lambda) - aF_1(\lambda+\alpha) + ap^{\lambda} F_N(\lambda+\alpha) = 0. \end{aligned} \tag{3}$$

Note that the first row of Eq. (3) is recursive with respect to $k$, which indicates that $F_{k-1}$ can be represented by $F_k$ for each $2 \le k \le N$. Hence $F_k$ can be represented by $F_N$ as

$$F_k(\lambda) = \sum_{l=0}^{N-k} C_{N-k,l} \left(-\frac{A}{\alpha N}\right)^l (\lambda-\alpha)\cdots(\lambda-l\alpha) F_N(\lambda-l\alpha), \quad 1 \le k \le N. \tag{4}$$

Summing over $k$ in the above equation yields

$$F(\lambda) = \sum_{k=1}^{N} F_k(\lambda) = \sum_{k=0}^{N-1}\sum_{l=0}^{k} C_{k,l} \left(-\frac{A}{\alpha N}\right)^l (\lambda-\alpha)\cdots(\lambda-l\alpha) F_N(\lambda-l\alpha). \tag{5}$$

Inserting Eq. (4) into the second row of Eq. (3) shows that $F_N$ is the solution to the following functional equation:

$$p^{\lambda-\alpha} F_N(\lambda) = \sum_{l=0}^{N} C_{N,l} \left(-\frac{A}{\alpha N}\right)^l (\lambda-\alpha)\cdots(\lambda-l\alpha) F_N(\lambda-l\alpha). \tag{6}$$

Complex computations show the solution of this functional equation can be computed explicitly as follows.

**Theorem 1.** The solution of Eq. (6) is given by

$$F_N(\lambda) = -\frac{KA}{N}\Gamma\left(1-\frac{\lambda}{\alpha}\right)^{-1} \int_0^\infty u^{-\frac{\lambda}{\alpha}} \prod_{n=0}^{\infty} a_N(p^{\alpha n}u)du, \tag{7}$$

where

$$a_N(u) = \left(1+\frac{Au}{N}\right)^{-N}$$

is a function of $u$ and

$$K = \left[\int_0^\infty \frac{1}{u}(a_N(u)^{-1}-1) \prod_{n=0}^{\infty} a_N(p^{\alpha n}u)du\right]^{-1}$$

is a normalization constant.

*Proof.* The proof of this theorem is highly nontrivial and will be given in Section 5. □

Inserting Eq. (7) into Eq. (5) gives the following explicit expression for the moment generating function:

$$F(\lambda) = K\sum_{k=0}^{N-1}\sum_{l=0}^{k} C_{k,l} \left(\frac{A}{N}\right)^{l+1} \Gamma\left(1-\frac{\lambda}{\alpha}\right)^{-1} \int_0^\infty u^{l-\frac{\lambda}{\alpha}} \prod_{n=0}^{\infty} a_N(p^{\alpha n}u)du. \tag{8}$$

For convenience, we also introduction the characteristic function

$$G(\lambda) = \int_{-\infty}^{\infty} p(x) e^{i\lambda x} dx,$$

which is nothing but the inverse Fourier transform of $p(x)$. Clearly, the moment generating function and the characteristic function are related by $G(\lambda) = F(i\lambda)$. Thus we finally obtain the following explicit expression of the characteristic function:

$$G(\lambda) = K \sum_{k=0}^{N-1} \sum_{l=0}^{k} C_{k,l} \left(\frac{A}{N}\right)^{l+1} \Gamma\left(1 - \frac{i\lambda}{\alpha}\right)^{-1} \int_0^{\infty} u^{l-\frac{i\lambda}{\alpha}} \prod_{n=0}^{\infty} a_N(p^{\alpha n} u) du. \tag{9}$$

Since the Fourier transform and inverse Fourier transform are inverses of each other, taking the Fourier transform of the characteristic function $G(\lambda)$ yields the probability density $p(x)$ of the logarithmic cell size. Finally, the probability density of the original cell size $y = e^x$ is given by

$$\tilde{p}(y) = \frac{1}{y} p(\log y).$$

#### 2.2 Case of small cell cycle duration variability

We next focus on the special case where cell cycle duration variability is very small, i.e. $N \gg 1$. In this limit, we have $a_N(u) = e^{-Au}$ and thus we have

$$\prod_{n=0}^{\infty} a_N(p^{\alpha n} u) = \prod_{n=0}^{\infty} e^{-Ap^{\alpha n} u} = e^{-\frac{Au}{1-p^{\alpha}}}.$$

This implies that

$$\int_0^{\infty} u^{l-\frac{\lambda}{\alpha}} \prod_{n=0}^{\infty} a_N(p^{\alpha n} u) du = \int_0^{\infty} u^{l-\frac{\lambda}{\alpha}} e^{-\frac{Au}{1-p^{\alpha}}} du = \Gamma\left(l + 1 - \frac{\lambda}{\alpha}\right) \left(\frac{A}{1-p^{\alpha}}\right)^{\frac{\lambda}{\alpha}-l-1}.$$

Inserting this equation into Eq. (8) yields

$$F(\lambda) = K \sum_{k=0}^{N-1} \sum_{l=0}^{k} C_{k,l} \left(\frac{A}{N}\right)^{l+1} \Gamma\left(1 - \frac{\lambda}{\alpha}\right)^{-1} \Gamma\left(l + 1 - \frac{\lambda}{\alpha}\right) \left(\frac{A}{1-p^{\alpha}}\right)^{\frac{\lambda}{\alpha}-l-1}.$$

By virtue of the fact that

$$\frac{\Gamma(l+x)}{\Gamma(x)} = x(x+1)\cdots(x+l-1) = (x)_l,$$

we obtain

$$F(\lambda) = K \left(\frac{A}{1-p^{\alpha}}\right)^{\frac{\lambda}{\alpha}} \sum_{k=0}^{N-1} \sum_{l=0}^{k} C_{k,l} \left(\frac{1-p^{\alpha}}{N}\right)^{l+1} \left(1 - \frac{\lambda}{\alpha}\right)_l,$$

where $(x)_l = x(x+1)\cdots(x+l-1)$ is the Pochhammer symbol. Moreover, using the hockey-stick identity

$$\sum_{k=1}^{N-1} C_{k,l} = C_{N,l+1},$$

we obtain

$$\begin{aligned} F(\lambda) &= K\left(\frac{A}{1-p^\alpha}\right)^{\frac{\lambda}{\alpha}} \sum_{l=0}^{N-1}\sum_{k=l}^{N-1} C_{k,l}\left(\frac{1-p^\alpha}{N}\right)^{l+1}\left(1-\frac{\lambda}{\alpha}\right)_l \\ &= K\left(\frac{A}{1-p^\alpha}\right)^{\frac{\lambda}{\alpha}} \sum_{l=0}^{N-1} C_{N,l+1}\left(\frac{1-p^\alpha}{N}\right)^{l+1}\left(1-\frac{\lambda}{\alpha}\right)_l \\ &= \frac{K}{N}\left(\frac{A}{1-p^\alpha}\right)^{\frac{\lambda}{\alpha}} \sum_{l=0}^{N-1} \frac{C_{N-1,l}}{l+1}\left(\frac{1-p^\alpha}{N}\right)^{l+1}\left(1-\frac{\lambda}{\alpha}\right)_l. \end{aligned}$$

In the limit of $N \to \infty$, we have

$$\sum_{l=0}^{N-1} \frac{C_{N-1,l}}{l+1}\left(\frac{1-p^\alpha}{N}\right)^{l}\left(1-\frac{\lambda}{\alpha}\right)_l = \frac{\alpha(1-p^\lambda)}{\lambda(1-p^\alpha)},$$

This identity, together with the fact that $F(0) = 1$, finally shows that

$$F(\lambda) = \frac{1-p^\lambda}{-\lambda\log p}\left(\frac{A}{1-p^\alpha}\right)^{\frac{\lambda}{\alpha}}. \tag{10}$$

Direct computation shows that $F(\lambda)$ can be rewritten as

$$F(\lambda) = \frac{\bar{V}_d^\lambda - \bar{V}_b^\lambda}{(\log\bar{V}_d - \log\bar{V}_b)\lambda},$$

where

$$\bar{V}_b = p\left(\frac{A}{1-p^\alpha}\right)^{\frac{1}{\alpha}}, \quad \bar{V}_d = \left(\frac{A}{1-p^\alpha}\right)^{\frac{1}{\alpha}}.$$

Replacing $\lambda$ by $i\lambda$ in the above equation gives the characteristic function

$$G(\lambda) = \frac{\bar{V}_d^{i\lambda} - \bar{V}_b^{i\lambda}}{(\log\bar{V}_d - \log\bar{V}_b)i\lambda}.$$

Taking the Fourier transform of the characteristic function shows that logarithmic cell size has the uniform distribution

$$p(x) = \frac{1}{\log\bar{V}_d - \log\bar{V}_b} I_{[\log\bar{V}_b,\log\bar{V}_d]}(x),$$

and thus the original cell size $y = e^x$ has the following distribution:

$$\tilde{p}(y) = \frac{1}{y}p(\log y) = \frac{1}{(\log\bar{V}_d - \log\bar{V}_b)y} I_{[\bar{V}_b,\bar{V}_d]}(y), \tag{11}$$

where $I_A(x)$ is the indicator function which takes the value of 1 when $x \in A$ and takes the value of 0 otherwise.

#### 3 Cell cycle duration distribution

Let $V_b$ and $V_d$ denote the cell sizes at birth and at division in a particular generation, respectively, and let $T$ denote the corresponding cell cycle duration. Since the cell size growth exponentially, we have

$$V_d = V_b e^{gT}.$$

This shows that

$$T = \frac{1}{g}\log\left(\frac{V_d}{V_b}\right). \tag{12}$$

Recall that $V_d^\alpha - V_b^\alpha$ has an Erlang distribution with shape parameter $N$ and rate parameter $N/A$. Thus given that $V_b^\alpha = x$, the probability density function of $V_d^\alpha$ is given by

$$\mathbb{P}(V_d^\alpha = y | V_b^\alpha = x) = \frac{N^N}{A^N(N-1)!}(y-x)^{N-1}e^{-\frac{N}{A}(y-x)}, \quad y \geq x.$$

Thus given that $V_b^\alpha = x$, it follows from Eq. (12) that the probability density function of the cell cycle duration $T$ is given by

$$\mathbb{P}(T = t | V_b^\alpha = x) = \frac{\alpha g N^N}{A^N(N-1)!}x^N(e^{\alpha gt}-1)^{N-1}e^{\alpha gt - \frac{N}{A}x(e^{\alpha gt}-1)}.$$

We next compute the distribution of $V_b$. To this end, let $V_b(k)$ and $V_d(k)$ denote the cell sizes at birth and at division in the $k$th generation, respectively. Under the assumption of deterministic partitioning, we have $V_b(k+1) = pV_d(k)$ and thus we obtain the recursive equation

$$V_b^\alpha(k+1) = p^\alpha[V_b^\alpha(k) + \Delta_k], \quad k \geq 0,$$

where $\Delta_k = V_d^\alpha(k) - V_b^\alpha(k)$ is the generalized added size in the $k$th generation, which has an Erlang distribution with shape parameter $N$ and rate parameter $N/A$. Using the recursive equation repeatedly, we obtain

$$V_b^\alpha(k) = p^{k\alpha}V_b(0)^\alpha + p^{k\alpha}\Delta_0 + p^{(k-1)\alpha}\Delta_1 + \cdots + p^\alpha\Delta_{k-1}. \tag{13}$$

Since $\Delta_0, \Delta_1, \Delta_2, \cdots$ are i.i.d. Erlang distributed random variables with shape parameter $N$ and rate parameter $N/A$, the Laplace transform of $\Delta_n$ is given by

$$\mathbb{E}e^{-\lambda\Delta_n} = \left(1 + \frac{A\lambda}{N}\right)^{-N} = a_N(\lambda).$$

It thus follows from (13) and the independence of $V_b(0), \Delta_0, \Delta_1, \Delta_2, \cdots$ that

$$\mathbb{E}e^{-\lambda V_b^\alpha(k)} = \mathbb{E}e^{-\lambda p^{k\alpha}V_b(0)^\alpha}\prod_{n=1}^{k}a_N(p^{n\alpha}\lambda). \tag{14}$$

Since the distribution of $V_b(k)$ converges to the steady-state distribution of the birth size as $k \to \infty$, taking $k \to \infty$ in Eq. (14) shows that the Laplace transform of $V_b^\alpha$ is given by

$$\mathbb{E}e^{-\lambda V_b^\alpha} = \prod_{n=1}^{\infty}a_N(p^{\alpha n}u) = \prod_{n=1}^{\infty}\left(1 + \frac{Ap^{\alpha n}\lambda}{N}\right)^{-N}. \tag{15}$$

Taking the inverse Laplace transform gives the probability density function of $V_b^\alpha$, from which is the probability density function of $V_b$ can be obtained. Finally, the distribution of the cell cycle duration $T$ is given by

$$\mathbb{P}(T = t) = \int_0^\infty \mathbb{P}(T = t | V_b^\alpha = x)\mathbb{P}(V_b^\alpha = x)dx. \tag{16}$$

A special case occurs when $\alpha$ is large (strong cell-size control) or when $p$ is small (smaller daughter tracking). Under the large $\alpha$ or small $p$ approximation, the term $p^{\alpha n}$ is negligible for $n \geq 2$ and it suffices to keep only the first term in the infinite product given in Eq. (15). In this case, the laplace transform of $V_b^\alpha$ reduces to

$$\mathbb{E}e^{-\lambda V_b^\alpha} \approx \left(1+\frac{Ap^\alpha\lambda}{N}\right)^{-N}.$$

Taking the inverse Laplace transform gives the birth size distribution

$$\mathbb{P}(V_b = x) = \frac{N^N x^{\alpha N-1}e^{-\frac{N}{Ap^\alpha}x^\alpha}}{(N-1)!A^N p^{\alpha N}}.$$

Inserting this equation into Eq. (16) gives the doubling time distribution

$$\mathbb{P}(T=t) = \frac{\alpha g(2N-1)!}{p^{\alpha N}[(N-1)!]^2}\cdot\frac{e^{\alpha gt}(e^{\alpha gt}-1)^{N-1}}{(p^{-\alpha}+e^{\alpha gt}-1)^{2N}}.$$

### 4 Cell size distribution under stochastic partitioning

We next focus on the case of stochastic partitioning at cell division. In this case, Eq. (4) in the main text can be converted to the following differential equations satisfied by the moment generating function:

$$\begin{aligned}\partial_t F_k(\lambda) &= g\lambda F_k(\lambda) - aF_k(\lambda+\alpha) + aF_{k-1}(\lambda+\alpha), \quad 2\le k\le N,\\ \partial_t F_1(\lambda) &= g\lambda F_1(\lambda) - aF_1(\lambda+\alpha) + a\hat{\mu}(\lambda)F_N(\lambda+\alpha),\end{aligned}$$

where

$$\hat{\mu}(\lambda) = \int_0^\infty \mu(y)e^{-\lambda y}dy = \int_0^1 f(x)x^\lambda dx$$

is the Laplace transform of $\mu(y)$. At the steady state, in analogy to the derivation of Eq. (5), we obtain

$$F(\lambda) = \sum_{k=0}^{N-1}\sum_{l=0}^{k} C_{k,l}\left(-\frac{A}{\alpha N}\right)^l(\lambda-\alpha)\cdots(\lambda-l\alpha)F_N(\lambda-l\alpha), \tag{17}$$

where $F_N$ is the solution to the following functional equation:

$$\hat{\mu}(\lambda-\alpha)F_N(\lambda) = \sum_{l=0}^{N} C_{N,l}\left(-\frac{A}{\alpha N}\right)^l(\lambda-\alpha)\cdots(\lambda-l\alpha)F_N(\lambda-l\alpha). \tag{18}$$

To proceed, we defined a new function

$$p(\lambda) = \left(\int_0^1 f(x)x^{\lambda-\alpha}dx\right)^{\frac{1}{\lambda-\alpha}}.$$

With this notation, Eq. (18) can be rewritten as

$$p(\lambda)^{\lambda-\alpha}F_N(\lambda) = \sum_{l=0}^{N} C_{N,l}\left(-\frac{A}{\alpha N}\right)^l(\lambda-\alpha)\cdots(\lambda-l\alpha)F_N(\lambda-l\alpha). \tag{19}$$

Comparing Eqs. (7) and (19), we can see that when the noise in partitioning is small, the solution of the above functional equation is approximately given by

$$F_N(\lambda) = -\frac{KA}{N}\Gamma\left(1-\frac{\lambda}{\alpha}\right)^{-1}\int_0^\infty u^{-\frac{\lambda}{\alpha}}\prod_{n=0}^{\infty}a_N(p(\lambda)^{\alpha n}u)du,$$

where the constant $K$ is given by

$$K = \left[\int_0^\infty \frac{1}{u}(a_N(u)^{-1}-1)\prod_{n=0}^{\infty} a_N(p(0)^{\alpha n}u)du\right]^{-1},$$

with $p(0)$ being the limit of $p(\lambda)$ as $\lambda \to 0$. Inserting this equation into Eq. (17) shows that the moment generating function is given by

$$F(\lambda) = K\sum_{k=0}^{N-1}\sum_{l=0}^{k} C_{k,l}\left(\frac{A}{N}\right)^{l+1}\Gamma\left(1-\frac{\lambda}{\alpha}\right)^{-1}\int_0^\infty u^{l-\frac{\lambda}{\alpha}}\prod_{n=0}^{\infty} a_N(p(\lambda)^{\alpha n}u)du.$$

Replacing the variable $\lambda$ in the the moment generating function by $i\lambda$ yields the characteristic function

$$G(\lambda) = K\sum_{k=0}^{N-1}\sum_{l=0}^{k} C_{k,l}\left(\frac{A}{N}\right)^{l+1}\Gamma\left(1-\frac{i\lambda}{\alpha}\right)^{-1}\int_0^\infty u^{l-\frac{i\lambda}{\alpha}}\prod_{n=0}^{\infty} a_N(p(i\lambda)^{\alpha n}u)du.$$

Taking the Fourier transform of the characteristic function gives the the probability density $p(x)$ of the logarithmic cell size. Then the probability density of the original cell size $y = e^x$ is given by

$$\tilde{p}(y) = \frac{1}{y}p(\log y).$$

### 5 Proof of Theorem 1

Here we shall give the detailed proof of Theorem 1 in Section 2. To this end, we must revisit the cell size distribution for the adder strategy.

#### 5.1 Cell size distribution for the adder strategy

For the adder strategy ($\alpha = 1$), we can compute the cell size distribution in an alternative way. For simplicity, we consider the case of deterministic partitioning, i.e. $f(z) = \delta(z-p)$. In this case, Eq. (3) in the main text reduces to

$$\begin{aligned}\partial_t\tilde{p}_k(y) &= -\partial_y[gy\tilde{p}_k(y)] + ay\tilde{p}_{k-1}(y) - ay\tilde{p}_k(y), \quad 2\le k\le N,\\ \partial_t\tilde{p}_1(y) &= -\partial_y[gy\tilde{p}_1(y)] + \frac{ay}{p^2}\tilde{p}_N\left(\frac{y}{p}\right) - ay\tilde{p}_1(y).\end{aligned} \tag{20}$$

To proceed, for each cell cycle stage $k$, we introduce the Laplace transform

$$H_k(\lambda) = \int_0^\infty \tilde{p}_k(y)e^{-\lambda y}dy, \quad H(\lambda) = \int_0^\infty \tilde{p}(y)e^{-\lambda y}dy,$$

where $\tilde{p}(y) = \sum_{k=1}^N \tilde{p}_k(y)$ is the probability density function of the cell size. Then Eq. (20) can be converted to the following differential equations:

$$\begin{aligned}\partial_t H_k(\lambda) &= (g\lambda + a)\partial_\lambda H_k(\lambda) - a\partial_\lambda H_{k-1}(\lambda), \quad 2\le k\le N,\\ \partial_t H_1(\lambda) &= (g\lambda + a)\partial_\lambda H_1(\lambda) - a\partial_\lambda H_N(p\lambda).\end{aligned}$$

At the steady state, we have

$$\begin{aligned}(g\lambda + a)H_k'(\lambda) - aH_{k-1}'(\lambda) &= 0, \quad 2\le k\le N,\\ (g\lambda + a)H_1'(\lambda) - aH_N'(p\lambda) &= 0.\end{aligned} \tag{21}$$

Note that the first row of Eq. (21) is recursive with respect to $k$, which indicates that $H'_{k-1}$ can be represented by $H'_k$ for each $2 \leq k \leq N$. Hence $H'_k$ can be represented by $H'_N$ as

$$H'_k(\lambda) = \left(1 + \frac{A'\lambda}{N}\right)^{N-k} H'_N(\lambda), \quad 1 \leq k \leq N, \tag{22}$$

where $A' = Ng/a$. Inserting this equation into the second row of Eq. (21) shows that $H'_N$ is the solution to the following functional equation:

$$H'_N(\lambda) = \left(1 + \frac{A'\lambda}{N}\right)^{-N} H'_N(p\lambda) = a_N(\lambda) H'_N(p\lambda).$$

Using the above equation repeatedly yields

$$H'_N(\lambda) = H'_N(0) \prod_{n=1}^{\infty} a_N(p^n \lambda). \tag{23}$$

Inserting the above equation into Eq. (22) and summing over $k$, we obtain

$$H'(\lambda) = \sum_{k=1}^{N} H'_k(\lambda) = \frac{N}{A'} H'_N(0)(a_N(\lambda)^{-1} - 1) \prod_{n=1}^{\infty} a_N(p^n \lambda).$$

Since $H(0) = 1$, integrating the above equation yields

$$H(\lambda) = 1 + \frac{N}{A'} H'_N(0) \int_0^{\lambda} \frac{1}{u}(a_N(u)^{-1} - 1) \prod_{n=0}^{\infty} a_N(p^n u) du.$$

Finally, using the fact that $H(\infty) = 0$, we obtain

$$H(\lambda) = 1 - K \int_0^{\lambda} \frac{1}{u}(a_N(u)^{-1} - 1) \prod_{n=0}^{\infty} a_N(p^n u) du,$$

where

$$K = -\frac{N}{A'} H'_N(0) = \left[\int_0^{\infty} \frac{1}{u}(a_N(u)^{-1} - 1) \prod_{n=0}^{\infty} a_N(p^n u) du\right]^{-1}.$$

is a normalization constant. Taking the inverse Laplace transform gives the cell size distribution.

#### 5.2 Detailed proof

We are now in a position to prove Theorem 1.

*Proof.* For simplicity, we first focus on the adder strategy ($\alpha = 1$). Recall that the probability density functions at a certain stage for the original cell size and the logarithmic cell size are related by

$$\tilde{p}_k(y) = \frac{1}{y} p_k(\log y).$$

Using the change of variables formula, we obtain

$$F_N(\lambda) = \int_{-\infty}^{\infty} p_k(x) e^{\lambda x} dx = \int_0^{\infty} \tilde{p}_k(y) y^{\lambda} dy.$$

Straightforward computations shows that

$$\int_0^\infty x^{\lambda-1}e^{-yx}dx = \Gamma(\lambda)y^{-\lambda}.$$

Combining the above two equations shows that

$$\begin{aligned}F_N(-\lambda) &= \int_0^\infty \tilde{p}_k(y)y^{-\lambda}dy \\ &= \Gamma(\lambda)^{-1}\int_0^\infty \tilde{p}_k(y)dy\int_0^\infty x^{\lambda-1}e^{-yx}dx \\ &= \Gamma(\lambda)^{-1}\int_0^\infty x^{\lambda-1}dx\int_0^\infty \tilde{p}_k(y)e^{-xy}dy \\ &= \Gamma(\lambda)^{-1}\int_0^\infty H_k(x)x^{\lambda-1}dx.\end{aligned}$$

It follows from Eq. (23) that

$$H_N(x) = -\frac{KA'}{N}\int_x^\infty \prod_{n=0}^\infty a_N(p^n u)du.$$

Thus we obtain

$$\begin{aligned}F_N(-\lambda) &= -\frac{KA'}{N}\Gamma(\lambda)^{-1}\int_0^\infty x^{\lambda-1}dx\int_x^\infty \prod_{n=0}^\infty a_N(p^n u)du \\ &= -\frac{KA'}{N}\Gamma(\lambda)^{-1}\int_0^\infty \prod_{n=0}^\infty a_N(p^n u)du\int_0^u x^{\lambda-1}dx \\ &= -\frac{KA'}{N}\Gamma(\lambda+1)^{-1}\int_0^\infty u^\lambda \prod_{n=0}^\infty a_N(p^n u)du.\end{aligned} \tag{24}$$

It follows from Eq. (6) that $F_N(\lambda)$ is the solution to the functional equation

$$p^{\lambda-1}F_N(\lambda) = \sum_{l=0}^N C_{N,l}\left(-\frac{A'}{N}\right)^l (\lambda-1)\cdots(\lambda-l)F_N(\lambda-l). \tag{25}$$

Thus the solution of the above functional equation can be computed explicitly as

$$F_N(\lambda) = -\frac{KA'}{N}\Gamma(1-\lambda)^{-1}\int_0^\infty u^{-\lambda}\prod_{n=0}^\infty a_N(p^n u)du. \tag{26}$$

We next focus on the case of a general size control strategy (arbitrary $\alpha$). In this case, it follows from Eq. (6) that $F_N(\lambda)$ is the solution to the functional equation

$$p^{\lambda-\alpha}F_N(\lambda) = \sum_{l=0}^N C_{N,l}\left(-\frac{A}{\alpha N}\right)^l (\lambda-\alpha)\cdots(\lambda-\alpha)F_N(\lambda-\alpha). \tag{27}$$

Comparing the functional forms of Eqs. (24) and (27), it is easy to see that the solution to Eq. (27) can be obtain from the function given in Eq. by replacing $\lambda$ by $\lambda/\alpha$, replacing $A'$ by $A = \alpha A'$, and replacing $p$ by $p^\alpha$. Thus we finally obtain

$$F_N(\lambda) = -\frac{KA}{N}\Gamma\left(1-\frac{\lambda}{\alpha}\right)^{-1}\int_0^\infty u^{-\frac{\lambda}{\alpha}}\prod_{n=0}^\infty a_N(p^{\alpha n}u)du. \tag{28}$$

This gives the desired result. □
